# HIV-1 can undergo Env-independent retrotransposon-like activity

**DOI:** 10.64898/2026.04.21.719790

**Authors:** Sojiro Matsumura, Wright Andrews Ofotsu Amesimeku, Samiul Alam Rajib, Nami Monde, Hiroyuki Sasaki, Hiromi Terasawa, Md. Jakir Hossain, Tomohiro Sawa, Yosuke Maeda, Yorifumi Satou, Kazuaki Monde

## Abstract

Retrovirus replication requires coordinated interplay between viral proteins and host cellular machinery, including reverse transcription of the viral RNA genome into DNA and its subsequent integration into the host genomes. SOX2-dependent retrotransposon dynamics have been reported for endogenous retrovirus HERV-K; however, whether a similar intracellular pathway exists for exogenous retroviruses remains unclear. To address whether infection-independent intracellular reverse transcription and partial integration can occur, that is, whether retrovirus can exhibit retrotransposon-like activity, we utilized HeLa and 293T cells, which express no HIV-1 entry receptors. We engineered an Env-deficient HIV-1 NL4-3-based reporter encoding a reverse-oriented, intron-disrupted nanoluciferase cassette that becomes expressible only after splicing followed by reverse transcription. We found that reporter activation depends on reverse transcriptase and protease activities. While integrase is dispensable for early expression, it is essential for long-term maintenance of the nanoluciferase signal. Integration site mapping using next-generation sequencing further confirmed that stable reporter activity requires integrase-dependent proviral insertion. Functional analysis of Gag revealed that membrane binding, multimerization, and budding are prerequisite steps for reporter activation. Concentrated virus preparations from culture supernatants failed to activate the reporter in 293T cells, ruling out a role for reinfection. Electron and confocal microscopy suggested that Gag or viral particles traffic through endosomal compartments. Furthermore, inhibition of dynamin- and clathrin-dependent endocytic pathways reduced reporter activity, indicating that these pathways contribute to efficient reporter activity.

Collectively, these finding support the conclusion that HIV-1 can undergo intracellular reverse transcription and partial integration in an infection-independent manner, prompting a reconsideration of the boundary between exogenous retroviruses and endogenous retroelements.

**Author summary:** Env-independent, infection-independent intracellular reverse transcription and integration in HIV-1 may contribute to integration-site diversity within the same cell. More broadly, this phenomenon suggests continuity between the retroviral life cycle and retrotransposition dynamics, therefore informing our understanding of host-virus coevolution, mechanisms of long-term persistence, and the redesign of therapeutic strategies targeting pre-integration steps.

## Introduction

Retroviruses replicate by reverse transcribing their RNA genomes into cDNA and integrating the cDNA into the host genome [1-3]. HIV-1, an exogenous retrovirus, generates new particles from transcripts derived from the provirus [4]. Particle assembly, maturation, and replication are coordinated by the Gag and Gag-Pol polyproteins. The Gag polyprotein forms a structural scaffold: the N-terminal matrix (MA) domain is recruited to the plasma membrane via PI(4,5)P_2_ binding; the capsid (CA) domain mediates lattice formation and participates in uncoating and nuclear import; and the nucleocapsid (NC) domain packages viral RNA through zinc finger motifs [5, 6]. Furthermore, the C-terminal p6 domain recruits the ESCRT machinery via its PTAP motif to facilitate viral budding [7]. Pol-encoded enzymes execute essential enzymatic functions after viral release: protease (PR) cleaves Gag/Gag-Pol precursors to enable maturation, reverse transcriptase (RT) synthesizes viral cDNA, and integrase (IN) processes and inserts viral DNA into the host genome, thereby completing the viral replication cycle [2-4].

HIV-1 assembly occurs at the plasma membrane, where the Gag polyprotein undergoes multimerization, recruiting viral RNA and envelope glycoproteins, and culminating in budding and release of virions from the cell surface. Virion release is driven by the host ESCRT machinery, after which the particles undergo protease-mediated maturation to become infectious. Notable, plasma membrane budding is considered the primary route of HIV-1 egress, especially in T lymphocytes [8, 9].

At the same time, HIV-1 particles have been reported to accumulate within intracellular late endosomes or multivesicular bodies (MVBs), particularly in macrophages and dendritic cells [10]. These particles may evade immune surveillance and are thought to be maintained as a reservoir within virus-containing compartments (VCCs) [11]. Moreover, VCCs are believed to be physically connected to the plasma membrane through narrow channels, allowing sequestration of virions while preserving potential routes for release upon cellular activation. Nevertheless, much remains unknown about the infectivity and dynamics of particles accumulated within intracellular VCCs [10-12].

Endogenous retroviruses, such as MusD and HERV-K, can undergo reverse transcription and integration in specific contexts, resembling retrotransposition [13-15]. Additionally, intracellular progression of reverse transcription and accumulation of unintegrated viral DNA have been documented for exogenous retroviruses, suggesting the potential existence of infection-independent replication intermediates [3, 16, 17]. It remains unclear whether this retrotransposon-like mechanism was acquired after endogenization or whether exogenous retroviruses inherently possessed such capabilities.

To address whether early HIV-1 replication steps can proceed independently of entry-mediated infection, we developed an Env-deficient HIV-1 NL4-3 system coupled with a splicing-, reverse transcription-, and integration-dependent nanoluciferase reporter. This system enables selective monitoring of intracellular replication events while excluding plasmid-derived transcription. Using genetic and pharmacologic perturbations, we delineate key Pol- and Gag-dependent requirements for reverse transcription and integration. Our findings demonstrate that HIV-1 can undergo infection-independent reverse transcription, revealing a hierarchical dependence on Gag-mediated assembly, protease-deriven maturation, and integrase-dependent chromosomal insertion, and uncovering mechanistic parallels with retrotransposon-like pathways. Together, this study establishes a framework for understanding intracellular HIV-1 replication and provides new insight into the origins of integration-site diversity observed in latent infection models [18].

## Results

### An Env-deleted, intron-dependent nanoluciferase reporter selectively monitors reverse transcription

To determine whether retrovirus can exhibit retrotransposon-like activity through intracellular, infection-independent reverse transcription and integration in HeLa and 293T cells, we designed a reporter system based on the “intron-insertion indicator” principle (Fig. 1A). Briefly, using an env-deleted HIV-1 (*NL4-3*) backbone, a reverse-oriented, intron-containing nanoluciferase reporter was inserted at the env deletion site, between the SV40 promoter and the poly(A) signal. This reporter is expressed only after splicing and reverse transcription [19]. To validate the design, we transfected 293T and HeLa cells with the construct. The wild-type (WT) reporter exhibited a time-dependent increase in nanoluciferase activity from days 1 to 6, whereas the ΔGagPol negative control remained at background levels (Figs. 1B and 1C), indicating that nanoluciferase expression depends on intracellular reverse transcription.

**Fig. 1.**
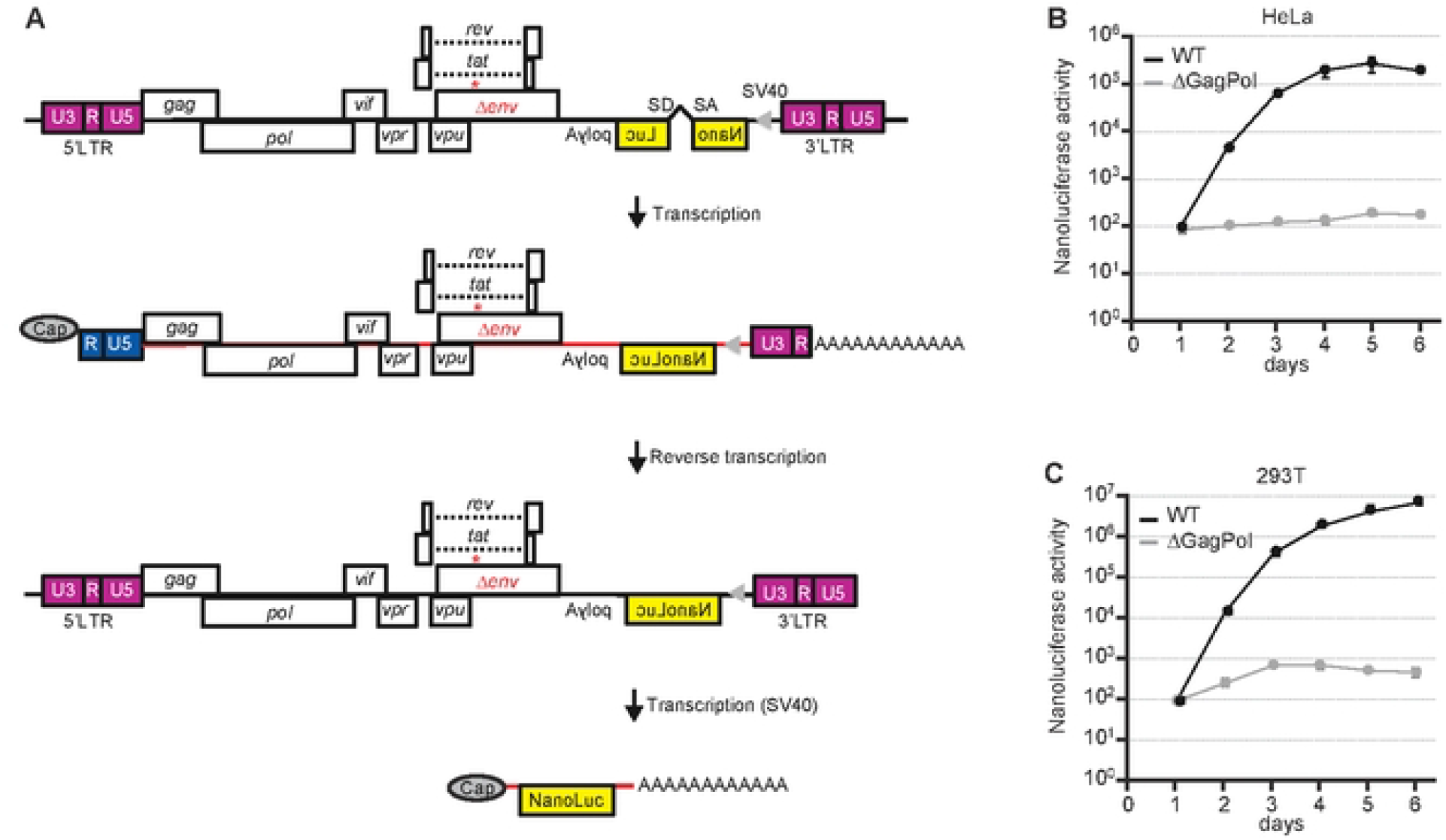
Assay design and initial evaluation. **(A)** Schematic of the Env-deleted NL4-3 construct containing a reverse-oriented Nluc reporter interrupted by an intron under an SV40 promoter, which becomes expressed only after reverse transcription and proper reorientation following delivery. **(B**-**C)** Time-course analysis of Nluc activity following plasmid transfection of HeLa (**B**) and 293T (**C**) cells using Lipofectamine. Nluc signals were monitored from Day 1 to Day 6 post-transfection.

### Reporter activation depends on reverse transcription, integration, and protease-mediated maturation

We next hypothesized that sustained reporter expression requires completion of downstream replication steps, including reverse transcription, integration and protease-mediated maturation. To investigate this, 293T cells were transfected with WT or *Pol* mutants lacking protease (Pro(-)), reverse transcriptase (RT(-)), or integrase (IN(-)), and nanoluciferase activity was monitored over seven days. Pro(-) and RT(-) mutants exhibited marked reduced nanoluciferase activity at all time points, whereas IN(-) showed activity comparable to WT during days 1-3 but declined from days 4-6 (Fig. 2A). These results are consistent with the canonical mechanism in which the processed of full-length linear vDNA generated by reverse transcription serve as substrates for integrase, leading to stable genomic integration [1, 3]. However, in the absence or inhibition of IN, unintegrated viral DNA species may transiently persist. These episomal DNAs can be degraded or diluted through cell division, and a fraction may circularize into relatively stable episomes [6, 17, 20]. Collectively, these reported fates provide a framework consistent with the time-dependent changes in reporter signal observed in our system. Western blot analysis of cell lysates and supernatants collected under identical conditions revealed comparable expression levels of the Gag precursor pr55 across samples (Fig. 2B) [21] , suggesting that the reductions in nanoluciferase activity observed in Fig. 2A reflect specific blocks in reverse transcription or maturation rather than differences in protein expression or transfection efficiency.

**Fig. 2.**
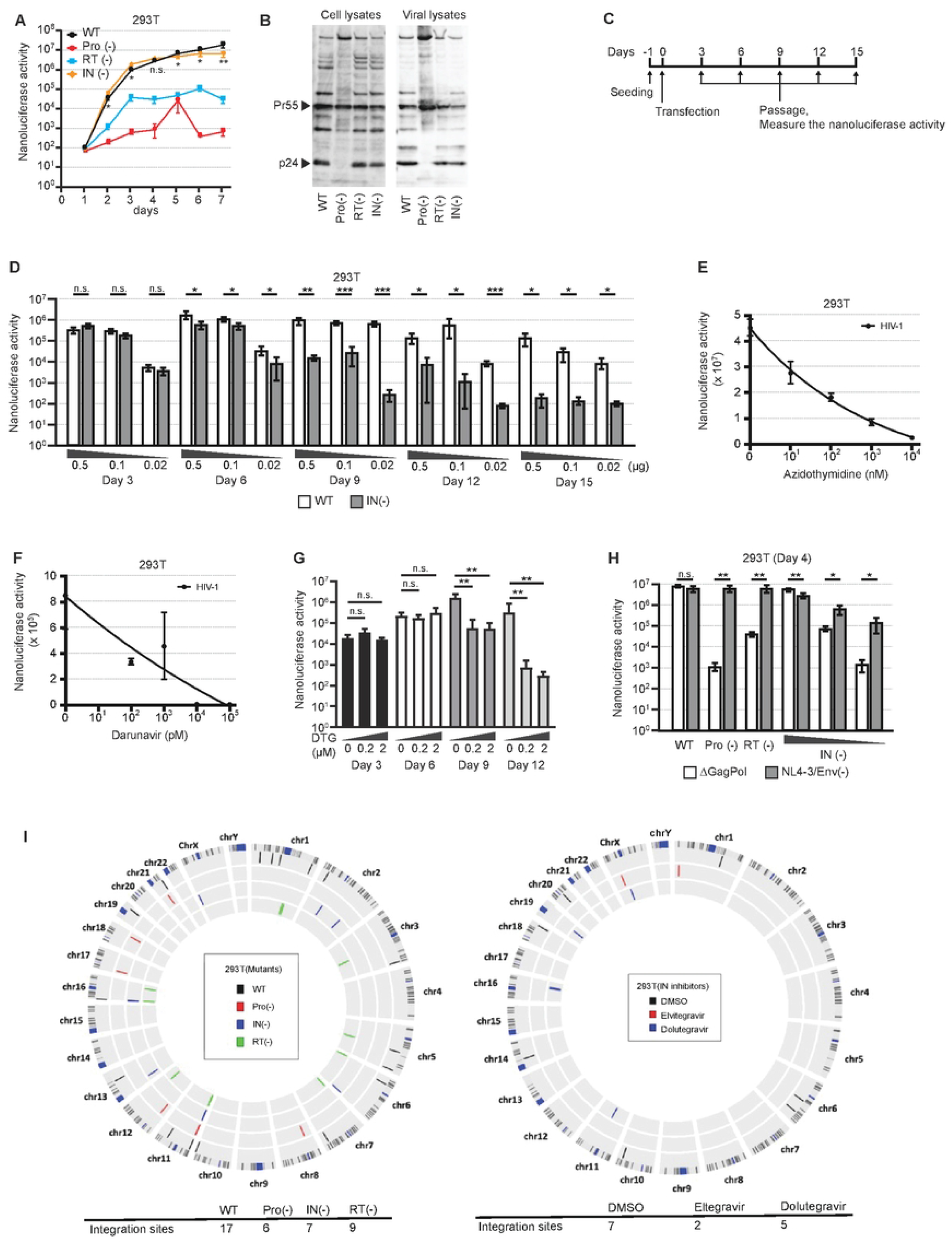
Pol dependencies and contribution of integration. **(A)** Nluc activity from Day 1 to Day 7 in 293T cells transfected with WT, Pro(-), RT(-), or IN(-) constructs. **(B)** Western blot analysis of cell and virus lysate under the same conditions. **(C** and **D)** Serial passaging of transfected cells (Days 3, 6, 9, 12, 15) using WT or IN(-) constructs at input DNA amounts of 0.5, 0.1, and 0.02 μg. **(E**-**G)** Nluc activity in the presence of AZT (**E**) or darunavir (**F**) or Doltegravir (DTG) (**G**). **(H)** Co-transfection of Pol mutants with ΔGagPol or NL4-3/Env(-). Nluc activity was measured on Day 4. **(I)** Bulk NGS analysis of integration sites, including Circos visualization and site counts. *Data are mean ± s.d. from three independent experiments.

To further investigate the functional consequences of integrase loss, we transfected WT and IN(-) constructs at various plasmid concentrations (0.5, 0.1 and 0.02 μg) in 293T cells and passaged every three days (Fig. 2C). Nanoluciferase outputs were similar on day 3 (Fig. 2D), indicating that early events driven directly by plasmid transcription occur efficiently even in the absence of integrase. However, significant differences emerged from day 6 onward: WT signals remained stable across passages, whereas IN(-) signals declined progressively. This difference reflects the inability of IN(-) particles to generate new proviruses once the initial plasmid-derived expression diminishes, demonstrating that integrase activity is dispensable for early expression but essential for maintaining replication competence over time. Hence, Fig. 2C captures the transition from transfection-driven expression to replication-driven propagation, identifying integrase as a key determinant of long-term viral fitness. Treatment with zidovudine (AZT: a reverse transcriptase inhibitor) [22], darunavir (a protease inhibitor) [23], dolutegravir (a integrase inhibitor) also produced dose- and time-dependent decreases in nanoluciferase activity (Fig. 2E-G), further supporting the dependencies on vDNA synthesis and protease-mediated maturation [24]. In trans-complementation assays, co-transfection with NL4-3/Env(-) restored nanoluciferase activity in Pol mutants (Fig. 2H), suggesting that the replication activities can be complemented in trans within the same cell [25]. Analysis of integration sites using next-generation sequencing (NGS) yielded variant counts of 17, 6, 7, and 9 for WT, Pro(-), IN(-), and RT(-), respectively; for WT treated with DMSO, elvitegravir [26], or dolutegravir [27], the counts were 7, 2, and 5 (Fig. 2I). Altogether, these results demonstrate that reporter activation depends on productive reverse transcription and protease processing, whereas integration is required for long-term maintenance of the signal.

### Gag-mediated membrane targeting and assembly are essential for reporter activation

We next examined whether Gag-mediated particle assembly influences retrotransposition. Gag membrane binding, multimerization-driven assembly, and recruitment of the ESCRT mechinery are essential for productive viral particle formation and release. To address this, we generated various mutants targeting distinct Gag domains. Specifically, ΔNC impairs RNA-dependent multimerization, selective packaging [28], and NC-dependent oligomerization/localization [29]. The 1GA mutation in the matrix domain of Gag reduces myristoylation-dependent membrane binding [30-32]. The 6A2T mutation, which also resides within the MA HBR, mediates PI(4,5)P_2_-dependent membrane binding and is critical for targeting Gag to the plasma membrane [33-34]. P99A, a mutation in the HIV-1 capsid, exhibits a “pre-budding” phenotype with defective membrane curvature formation [35, 36], whereas the WM184,185AA mutation disrupts CA-CTD dimerization and severely impairs assembly [37, 38]. A p6 domain mutation, PTAP(-), inhibits budding by blocking ESCRT recruitment (TSG101) [7].

We observed reductions in the nanoluciferase activity relative to WT for all mutants, while western blots confirmed preserved intracellular Gag expression (Pr55/p24) (Figs. 3A and 3B). These results indicate that the decrease in reporter activity is not due to altered expression levels or transfection efficiency, suggesting that Gag domain functions required for particle assembly at the plasma membrane are critical for retrotransposon activity.

**Fig. 3.**
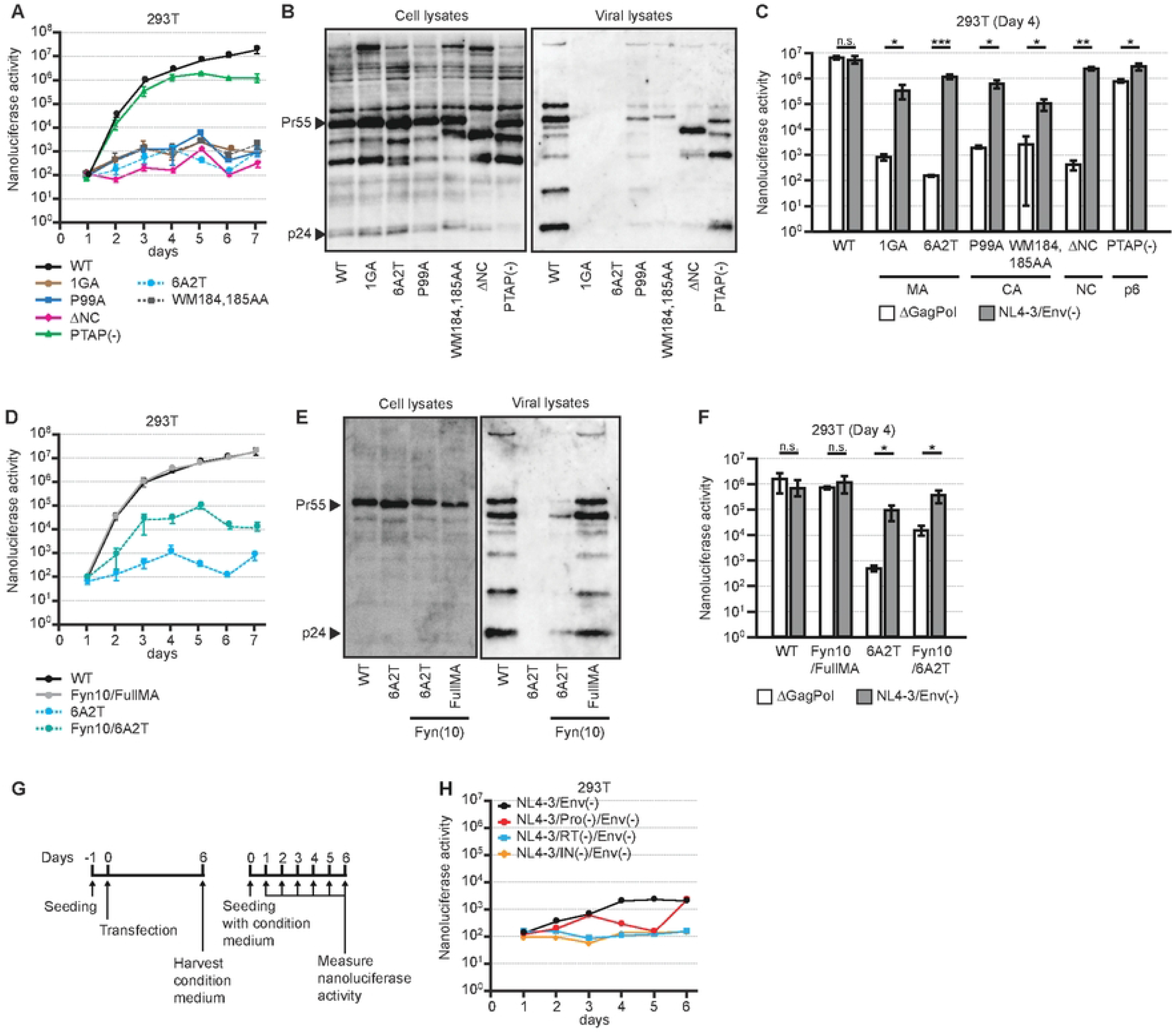
Gag functional requirements and complementation; exclusion of reinfection. **(A)** Nluc activity from day’s 1 to 7 in 293T cells transfected with WT or Gag mutants (ΔNC, 1GA, 6A2T, P99A, WM184,185AA, PTAP(-)). **(B)** Western blot analysis of cell and virus lysate of WT or Gag mutants as **(A). (C)** Complementation of each Gag mutant by co-transfection with ΔGagPol or NL4-3/Env(-). Nluc activity was measured on Day 4. **(D)** Time-course Nluc activity (Days 1-7) for 6A2T, Fyn10/6A2T, Fyn10/FullMA, and WT constructs. **(E)** Western blot analysis of cell and virus lysate for 6A2T, Fyn10/6A2T, Fyn10/FullMA, and WT constructs **(D). (F)** Complementation assays in which NL4-3/Env(-) was co-transfected with 6A2T or Fyn10/6A2T. Nluc activity was assayed on Day 4. **(G** and **H)** Reinfection control: concentrated supernatants were applied to naïve 293T cells, and Nluc activity was measured over Days 1–6. *Data are mean ± s.d. from three independent experiments.

Nanoluciferase activity was partially restored for each mutant when co-transfected with NL4-3/Env(-), relative to ΔGagPol (Fig. 3C), further indicating that Gag/Pol functions can be complemented in trans. Furthermore, the fusion of Fyn10, a palmitoylation-dependent non-specific membrane anchor, to the 6A2T mutant partially restored nanoluciferase activity compared to 6A2T alone, indicating that impaired membrane association contributes to the functional defect of this mutant. However, the Fyn10/6A2T construct did not reach WT levels, suggesting that simple membrane binding is insufficient to fully substitute for native MA-mediated membrane interactions. In contrast, appending Fyn10 to WT MA had no effect on activity, further demonstrating the importance of proper Gag targeting to the plasma membrane (Figs. 3D and E).

Functional complementation assays further demonstrated that co-transfection with NL4-3/Env(-) significantly enhanced activity for 6A2T and Fyn10/6A2T but not for WT or Fyn10/FullMA. Collectively, these results indicate that proper Gag-specific membrane targeting and assembly at the plasma membrane are essential for retrotransposon activity.

### Nanoluciferase activity arises independently of reinfection

We next examined whether nanoluciferase activity might result from secondary infection of neighboring cells by virions released from the plasma membrane, rather than from retrotransposition occurring within the originally transfected cells. To test this, viruses in the supernatants harvested at day 6 from WT- or Pol-mutant–transfected cells were concentrated 100-fold and applied to CD4-negative 293T cells. No increase in nanoluciferase activity was detected over days 1-6 (Figs. 3G and 3H), indicating that the reporter signal observed in earlier figures (Figs. 1-3) does not depend on reinfection of the neighboring cells but instead reflects Env-independent intracellular retrotransposon-like replication.

### Clathrin-mediated endocytosis drives intracellular trafficking of viral particles/RNPs

Because Gag is critical, we hypothesized that intracellularly accumulated viral particles or rebonucleoprotein (RNP)-like structures may influence retrotransposon-like replication. To visualize vRNA, a construct containing 24 × MS2 stem loop in RT and MS2-YFP were cotransfected into HeLa cells, and the localiation of Gag-mRFP and MS2-YFP was examined by confocal microscopy. On day 1, both Gag and vRNA were localized at the plasma membrane (Figs. 4A and 4B). By day 2, they accumulated not only at the plasma membrane but also within intracellular compartments (Fig. 4B). We next used transmission electron microscopy to determine whether virus-containing compartment (VCCs), as previously reported [39], are present in HIV-1-transfected HeLa cells. We found that mature particles containing cone-shaped cores were present within VCCs, even in the absence of Env and the CD4 receptor (Fig. 4C). These results suggest that virion can be internalized and trafficked through endosomes even in the absence of Env-receptor interactions.

**Fig. 4.**
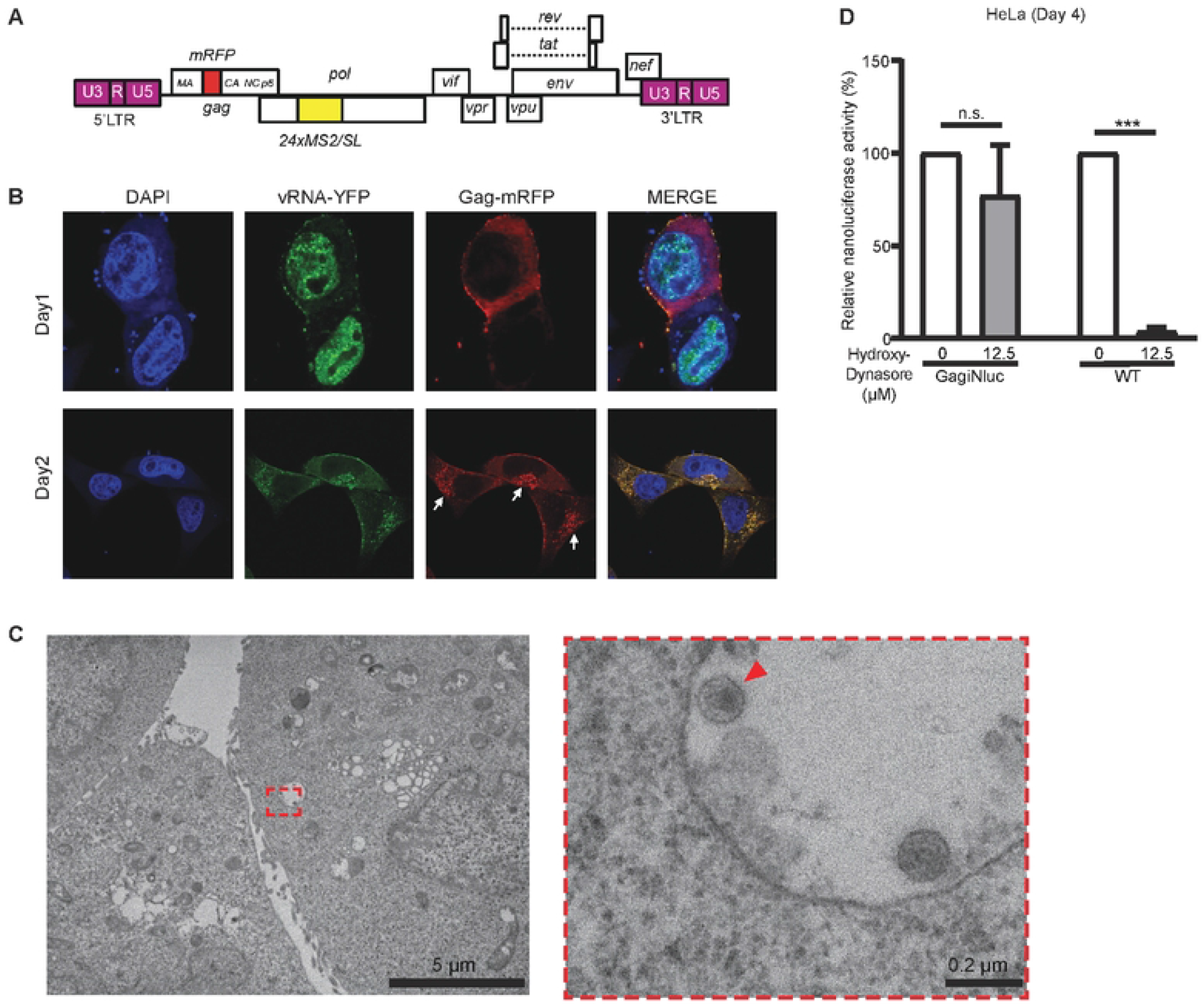
Intracellular dynamics visualized by transmission electron microscopy and confocal microscopy. **(A** and **B)** Confocal imaging of NL4-3 expressing mRFP-Gag with vRNA labeled using the 24×MS2/SL system (Construct as shown in **B**). Representative Day 1 and Day 2 images show vRNA (green), Gag (red), and nuclei (DAPI, blue). **(C)** Transmission electron microscopy (TEM) showing virus-like particles within endosomal compartments. Scale-200 nm. **(D)** Nanoluciferase activity in cells transfected with either Gag-iNluc (transfection-efficiency control) or WT, treated with DMSO or hydroxy-dynasore. *Data are mean ± s.d. from three independent experiments.

Because dynamin has been implicated in clathrin-mediated endocytosis, we next tested whether uptake pathways influence retrotransposon-like replication. Cells were treated with hydroxy-dynasore [40] to inhibit dynamin-mediated endocytosis, which resulted in a marked reduction in nanoluciferase activity (Fig. 4D). To ensure that hydroxy-dynasore did not affect transfection efficiency, we measured the nanoluciferase activity of Gag-iNLuc, a construct in which NLuc is inserted between the MA and CA domains of Gag, and observed no significant difference. These results indicate that clathrin-mediated endocytosis plays an important role in retrotransposon-like replication. Taken together, these finding suggest that clathrin-mediated endocytosis allows HIV-1 particles to accumulate in VCCs, after which some cores may re-enter the cytoplasm, undergo reverse transcription, and potentially integrate into the host genome.

## Discussion

In this study, we used an Env-deficient HIV-1 NL4-3 backbone carrying a reverse-oriented, intron-containing nanoluciferase reporter that becomes expressible only after reverse transcription to probe intracellular replication steps in CD4 receptor-negative 293T and HeLa cells. Using this system, we demonstrate that transmission-independent intracellular reverse transcription occurs and that sustained nanoluciferase activity over time depends on integrase-mediated integration. We further show that Gag assembly at the plasma membrane and particle accumulation within VCCs are important for intracellular retrotransposon-like replication. Because packaging of vRNA within the core is critical for HIV-1 reverse transcription and nuclear import of the viral genome, it is not surprising that core formation is also required for retrotransposon-like replication. However, receptor-independent viral accumulation in VCCs suggests a mechanism distinct from canonical viral entry.

We propose a working model in which transfected plasmid DNA initiates a cyclical pathway involving transcription, translation, Gag assembly and budding, endocytic re-internalization, reverse transcription, and ultimately integration. This pathway is capable of driving retrotransposon-like activity under specific conditions. In principle, substituting an already integrated provirus for the plasmid DNA yields a scenario in which intracellular genome copies could increase without classical infection, echoing intracellular retrotransposition described for HERV-K [13]. Supporting this idea, multiplicity of infection (MOI) is typically interpreted as the result of cells being infected by multiple viral particles. For example, studies of splenic germinal centers have reported rare HIV-positive cells containing clusters of related sequences and recombinants, interpreted as evidence for multiple infection, although the underlying mechanism could not be resolved because analyses were performed on bulk tissue [41]. Single-cell analyses of splenic lymphocytes have also identified an average of approximately three proviral copies per cell in vivo, indicating that multicopy cells do occur, although the mechanism was remains unclear [42]. In contrast, other single-cell studies report that most infected CD4^+^ T cells carry only one provirus, with only a small fraction showing evidence of multiple infection [43]. Moreover, this pathway may help explain how newly generated intracellular copies are amplified in an Env-independent manner and contribute to the integration-site heterogeneity observed in latent cell lines [18]. Although these observations are typically explained by simultaneous entry of multiple viral particles, the experimental approaches used do not clearly distinguish between classical multi-particle infection and potential intracellular genome amplification. Our Env-independent, retrotransposon-like mechanism therefore raises the possibility that intracellular replication contributes to a subset of cases previously attributed solely to multiple infection. Notably, our experimental system, which uses low receptor expression and an Env-deleted virus, was not designed to estimate in vivo rates of multiple infection. Fully distinguishing these mechanisms and their relative contributions will require single-cell resolution analyses in relevant primary cell types [41-43].

Consistent with previous reports, the effects of multiple Gag domain mutations on membrane targeting, assembly, and budding support the plasma membrane as the principal site of assembly site, even in cultured cells [9, 44, 45]. Furthermore, the observation of mature-like cores within endosomes and the sensitivity to dynamin inhibition suggest that clathrin-and dynamin-dependent uptake, followed by intracellular relocalization, contributes to the activity captured by our system [9, 20, 46, 47]. However, it remains unclear whether virions assembled at the plasma membrane are subsequently internalized via clathrin-dependent pathways and accumulate in VCCs, or whether Gag assembly intermedites are internalized and complete particle formation within late endosome [9, 44, 45]. Moreover, under Env-independent conditions, the route by which particle contents reach the cytosol remains unsolved. As a conceptual reference, single-particle imaging studies have shown that mature, Env-bearing virions fuse within endosomes and release their contents [48]. In our Env-independent system, however, the mechanism of cytosolic entry remains to be determined. A key limitation of this study is the restricted cell-type scope. Most experiments were performed in 293T and HeLa cells, and validation in physiologically relevant cell types (e.g., CD4^+^ T cells and macrophages) remains to be tested. HIV-1 entry exhibits cell type- and context-dependent plasticity; for example, clathrin-mediated uptake can operate in HeLa cells [47]. Single-particle analyses have demonstrated dynamin-dependent fusion within endosomes [47] and productive, endocytosis-dependent entry during cell-to-cell transmission [20]. These observations underscore the importance of validation in primary cells. At present, technical limitations preclude determining whether the Env-independent RT to (partial) integration framework demonstrated here also occurs in CD4^+^ T cells and monocyte-derived macrophages. For reservoir control, existing “shock and kill”[49] and “block and lock”[50] strategies could potentially be strengthened by explicitly suppressing intracellular RT and new de novo integration, for example by combining INSTIs, CA inhibitors, RT inhibitors, and LRAs or lock agents. Reduced integration-site diversity could serve as a potential marker of efficacy. More broadly, this mechanism blurs the distinction between exogenous retroviruses and endogenous retroelements, resembling Env-independent retrotransposition observed in HERV-K, and may refine our understanding of HIV-1 persistence, diversification, latency, and therapeutic opportunities.

## Acknowledgments

This work was supported by the Japan Agent for Medical Research and Development (AMED) Research Program on HIV/AIDS (JP24fk0410065h0001 and JP25fk0410065h0001) to K.M.; AMED Research Program on HIV/AIDS (JP25fk0410070, JP25fk0410071) to Y.S. AMED-multidisciplinary (JP23wm0325068) to Y.S., JST-ASPIRE Program (JP29jf0126018) to Y.S., JSPS KAKENHI Grants (JP23KK0292 and JP23K06561) to K.M., (JP25K02693) to Y.S. and the Takeda Science Foundation to K.M. The following reagent was obtained through the NIH HIV Reagent Program, Division of AIDS, NIAID, NIH: polyclonal Anti-Human Immunodeficiency virus Immune Globulin, Pooled Inactivated Human Sera, ARP-3957, contributed by NABI and National Heart Lung and Blood Institute (Dr. Luiz Barbosa), Zidovudine (AZT) (#HRP-3485) contributed by Creative Biolabs, Darunavir (ARP-11447) was obtained through BEI Resources. We thank Akira Ono for providing various Gag mutants.

## Material and methods

### Chemical compounds

Zidovudine (AZT; #HRP-3485) manufactured by TCI America, Inc, (Oregon, United States) was sourced from NIH AIDS Research and Preference Reagent Program, Division of AIDS at the National Institute of Allergy and Infectious Diseases. Darunavir (Brand name: Prezista; TMC 114), ARP-11447: DRV) was obtained through BEI Resources, NIAID, NIH, Elvitegravir (MedChem Express Company (MCE); Cat# HY-14740) and Dolutegravir (DTG) MedChemExpress(MCE); Cat# HY-13238) were obtained from the NIH AIDS Research and Reference Reagent Program, Division of AIDS, National Institute of Allergy and Infectious Diseases (NIAID).

### Plasmid

The HIV-1 molecular clone pNL4-3/Env(-) contains a frameshift mutation that disrupts Env expression [32]. Intron-disrupted nanoluciferase (inNanoluc) was designed as previously described [51] and was inserted between the SV40 promoter and the poly (A) signal near the 3′ LTR. Reverse transcriptase– and integrase-defective HIV-1 NL4-3 constructs were generated by site-directed mutagenesis, introducing the catalytic substitutions D110N in RT and D116N in integrase. The inNanoluc cassette encoded the SV40 early enhancer/promoter and SV40 late poly(A) signal positioned at an antisense orientation. The pNL4-3/Fyn (10) fullMA, pNL4-3/6A2T, pNL4-3/Fyn (10) 6A2T, pNL4-3/1GA, pNL4-3/P99A, pNL4-3/WM184,185AA, pNL4-3/PTAP(-), pNL4-3/24 × MS2/SL, and pMS2-YFP were elucidated in earlier studies [5, 31, 35, 36, 52] (a kind gift from A. Ono). pNL4-3/delNC is a kind gift from D. Ott [53]. The D25N protease-inactivating mutation was inserted into pNL4-3 to generate pNL4-3/Pro(-).

### Cells

HeLa and 293T cells were cultured in Dulbecco’s modified Eagle medium (Cat. No. D5796; DMEM; Sigma-Aldrich, Missouri, USA) supplemented with 5% fetal bovine serum (FBS; Cat. No. 173012; Sigma-Aldrich).

### Retrotransposition assay

293T cells were seeded into six-well plates at a density of 2×10^5^ cells/well. The cells were transfected with Lipofectamine 3000 reagent (Invitrogen), according to the manufacturer’s protocol. The cells were harvested 1 to 6 days after transfection, and the nanoluciferase activity in the cells was measured using the Nano-Glo luciferase assay reagent (Promega). In another case, azidothymidine and darunavir was treated immediately after transfection.

### Transmission electron microscopy analysis

HeLa cells were transfected with pNL4-3/Env(-)/inNanoluc. Viruses were collected by centrifugation at 13,200 × *g* for 1 h. After centrifugation, the supernatant was removed, leaving the viral pellet. The pelleted viruses were fixed with 2% glutaraldehyde (Cat. No. G010; GA; TAAB Laboratories Equipment, Aldermaston, England) and 1% osmium tetroxide (Cat. No. O018; TAAB Laboratories Equipment), followed by dehydration through a graded ethanol series (50%, 70%, 80%, 90%, 95%, and 99.5%). Samples were then embedded in Epon812 resin (Cat. No. T024; TAAB Laboratories Equipment). Ultrathin sections were prepared on copper grids (Cat. No. 2823; Nisshin EM, Tokyo, Japan), stained with Mayer’s hematoxylin solution (Cat. No. MHS16; Sigma-Aldrich) and lead citrate (Cat. No. 18-0875-2; Sigma-Aldrich), as described previously [54]. Stained sections were examined using a Hitachi 7600 transmission electron microscope (Hitachi High-Technologies, Tokyo, Japan) at 80 kV.

### Ligation-mediated PCR

Bulk viral integration sites were identified using ligation-mediated PCR (LM-PCR) followed by high-throughput sequencing, as previously described, with minor modifications [55]. Briefly, genomic DNA was isolated and one µg of the extracted DNA were fragmented by sonication using a picoruptor device (Diagenode) to produce fragments in the range of 300-400 base pairs. The DNA ends were repaired and the DNA linkers were added. The junction between the 3’ viral long terminal repeat (LTR) and host genomic DNA was amplified using linker-mediated PCR.

The first round of amplification employed the forward primer, HIV-1 - Bio3 (5’-GCTTGCCTTGAGTGCTCAAAGTAGTGT-3’) and reverse primer, Bio4 (5’-TCATGATCAATGGGACGATCA-3’) , followed by a nested PCR using HIV-1_P5Bio5 as Forward primer (5’-AATGATACGGCGACCACCGAGATCTACACGTGCCCGTCTGTTGTGT-3’) and PE-P7 as the Reverse primer (5’-CAAGCAGAAGACGGCATACGAGAT-3’). After the second round of nested PCR, the amplicons were quantified by qPCR using Illumina P5 and P7 primers.

Prepared libraries were sequenced as paired-end reads using the Illumina MiSeq platform. The sequencing primer targeting the viral 3’LTR , HIV-1-Seq Primer (5’-ATCCCTCAGACCTTTTAGTCAGTGTGG-3’), Adaptor Barcode Primer (5’-GATCGGAAGAGCGGTTCAGCAGGAAT-3’), and Adapter Sequencing Primer/ Read 2 Primer (5’-CGGTCTCGGCATTCCTGCTGAACCGCT-3’) were added to wells no. 12, 13, and 14, respectively, of the MiSeq cartridge during the sequencing run.

FASTQ files were processed for quality control and demultiplexing. To improve demultiplexing accuracy, an in-house script was used to retain reads with Phred quality scores >20 across all positions of the 8-bp index read.

Reads containing viral 3’LTR terminal sequences were identified, trimmed, and the remaining flanking host genomic sequences were mapped to the human reference genome (GRCh38/hg38) using the Burrows-Wheeler Aligner (BWA-MEM; v1.0) [56]. Reads with multiple alignments and PCR duplicates were removed using Samtools [56] and Picard [57] Integration sites were identified and quantified computationally, and selected sites were manually validated using the UCSC Genome Browser and BLAST against the NIH human genome database.

### Western blotting analysis

Cells and viruses were lysed with 1% Triton X lysis buffer (1% Triton X-100; Cat. No. 35501-02; Nacalai Tesque, Inc., Kyoto, Japan, 50 mM Tris-HCl, 300 mM NaCl, 10 mM iodoacetamide) supplemented with a protease inhibitor cocktail (Cat. No. 04693116001; Sigma-Aldrich). After lysis, SDS sample buffer was added. HIV-1 Gag was detected by immunoblotting using HIV-1 Immunoglobulin G (HIV-Ig: NIH AIDS reagent program). Horseradish peroxidase (HRP)-conjugated anti-human secondary antibody (Cat. No. 109-035-003; Jackson ImmunoResearch Laboratories, Pennsylvania, USA) was used. HRP signals were visualized using the ChemiDoc Touch Imaging System (BIO-RAD, California, USA).

### Confocal microscopy

HeLa cells were seeded onto collagen-coated 8-well chamber slides (Cat. No. 192-008; Watson Co., Ltd. Tokyo, Japan). Cells were co-transfected with pNL4-3/Gag-imRFP/24 × MS2/SL and pMS2/YFP using Lipofectamine 3000, as described above. At 16 h post-transfection, cells were fixed with 4% PFA for 30 min at 4 °C.

Permeabilization was carried out using 0.1% Triton X-100 for 2 min and then treated with 0.1 M glycine for 10 min, followed by blocking with 3% bovine serum albumin (BSA; Cat. No. 01863-48; Nacalai Tesque Inc.) for 30 min. Nuclei were stained with 4’,6-diamidino-2-phenylindole (DAPI) for 5 min and washed. Samples were then mounted with Dako Fluorescence Mounting Medium (Cat. No. S3023; Dako, Glostrup, Denmark), and imaged using Zeiss LSM 700 laser-scanning confocal microscopy.

### Hydroxy-Dynasore inhibition assay

HeLa cells were seeded in 12-well plates at 5 × 10^4^ cells/well and transfected with either the WT reporter plasmid or a Gag-iNluc control plasmid using Lipofectamine 3000 (Invitrogen) according to the manufacturer’s instructions. At 8 h post-transfection, the medium was removed and cells were washed twice with 1 × PBS, followed by addition of 1 mL/well fresh medium containing either vehicle (0.0625% DMSO, v/v) or Hydroxy-Dynasore (12.5 µM; from a 20 mM DMSO stock). Cells were harvested at 4 days post-transfection, and NanoLuc activity was measured using the Nano-Glo Luciferase Assay System (Promega).

### Statistical analysis

All statistical analyses were performed using GraphPad Prism9. One-way ANOVA with Sidak’s multiple comparison test and two-way ANOVA with Tukey’s multiple comparisons test were used as appropriate. Paired and unpaired student’s *t*-tests were also performed where indicated. Error bars represent the mean ± standard deviation from at least three independent experiments.

## References

1. Craigie R, Bushman FD. HIV DNA Integration. Csh Perspect Med. 2012;2(7). doi: ARTN a006890 10.1101/cshperspect.a006890. PubMed PMID: WOS:000314279100001.

2. Lesbats P, Engelman AN, Cherepanov P. Retroviral DNA Integration. Chem Rev. 2016;116(20):12730–57. Epub 20160520. doi: 10.1021/acs.chemrev.6b00125. PubMed PMID: 27198982; PubMed Central PMCID: PMCPMC5084067.

3. Hu WS, Hughes SH. HIV-1 Reverse Transcription. Csh Perspect Med. 2012;2(10). doi: ARTN a006882 10.1101/cshperspect.a006882. PubMed PMID: WOS:000314281700012.

4. Sundquist WI, Krausslich HG. HIV-1 assembly, budding, and maturation. Cold Spring Harb Perspect Med. 2012;2(7):a006924. doi: 10.1101/cshperspect.a006924. PubMed PMID: 22762019; PubMed Central PMCID: PMCPMC3385941.

5. Vineela Chukkapalli IBH, Vitaly Boyko, Wei-Shau Hu, Akira Ono. Interaction between the human immunodeficiency virus type 1 Gag matrix domain and phosphatidylinositol-(4,5)-bisphosphate is essential for efficient gag membrane binding. 2008:82(5):2405–17. doi: 10.1128/JVI.01614-07 PubMed Central PMCID: PMC PMC2258911.

6. Campbell EM, Hope TJ. HIV-1 capsid: the multifaceted key player in HIV-1 infection. Nature Reviews Microbiology. 2015;13(8):471–83. doi: 10.1038/nrmicro3503. PubMed PMID: WOS:000358000700006.

7. Garrus JE, von Schwedler UK, Pornillos OW, Morham SG, Zavitz KH, Wang HE, et al. Tsg101 and the vacuolar protein sorting pathway are essential for HIV-1 budding. Cell. 2001;107(1):55–65. doi: 10.1016/s0092-8674(01)00506-2. PubMed PMID: 11595185.

8. Freed EO. HIV-1 assembly, release and maturation. Nat Rev Microbiol. 2015;13(8):484–96. doi: 10.1038/nrmicro3490. PubMed PMID: WOS:000358000700007.

9. Jouvenet N, Neil SJD, Bess C, Johnson MC, Virgen CA, Simon SM, et al. Plasma membrane is the site of productive HIV-1 particle assembly. Plos Biol. 2006;4(12):2296–310. doi: ARTN e435 10.1371/journal.pbio.0040435. PubMed PMID: WOS:000242789100013.

10. Pelchen-Matthews A, Kramer B, Marsh M. Infectious HIV-1 assembles in late endosomes in primary macrophages. J Cell Biol. 2003;162(3):443–55. doi: 10.1083/jcb.200304008. PubMed PMID: WOS:000184667900010.

11. Koppensteiner H, Banning C, Schneider C, Hohenberg H, Schindler M. Macrophage Internal HIV-1 Is Protected from Neutralizing Antibodies. J Virol. 2012;86(5):2826–36. doi: 10.1128/Jvi.05915-11. PubMed PMID: WOS:000300536800040.

12. Bennett AE, Narayan K, Shi D, Hartnell LM, Gousset K, He HF, et al. Ion-Abrasion Scanning Electron Microscopy Reveals Surface-Connected Tubular Conduits in HIV-Infected Macrophages. Plos Pathog. 2009;5(9). doi: ARTN e1000591 10.1371/journal.ppat.1000591. PubMed PMID: WOS:000270804900007.

13. Monde K, Satou Y, Goto M, Uchiyama Y, Ito J, Kaitsuka T, et al. Movements of Ancient Human Endogenous Retroviruses Detected in SOX2-Expressing Cells. J Virol. 2022;96(9):e0035622. Epub 20220414. doi: 10.1128/jvi.00356-22. PubMed PMID: 35420440; PubMed Central PMCID: PMCPMC9093106.

14. Johnson WE. Endogenous Retroviruses in the Genomics Era. Annu Rev Virol. 2015;2(1):135–59. Epub 20150828. doi: 10.1146/annurev-virology-100114-054945. PubMed PMID: 26958910.

15. Ribet D, Dewannieux M, Heidmann T. An active murine transposon family pair: retrotransposition of “master” MusD copies and ETn trans-mobilization. Genome Res. 2004;14(11):2261–7. Epub 20041012. doi: 10.1101/gr.2924904. PubMed PMID: 15479948; PubMed Central PMCID: PMCPMC525684.

16. Geis FK, Goff SP. Unintegrated HIV-1 DNAs are loaded with core and linker histones and transcriptionally silenced. P Natl Acad Sci USA. 2019;116(47):23735–42. doi: 10.1073/pnas.1912638116. PubMed PMID: WOS:000498683000054.

17. Bukrinsky MI, Sharova N, Dempsey MP, Stanwick TL, Bukrinskaya AG, Haggerty S, et al. Active nuclear import of human immunodeficiency virus type 1 preintegration complexes. Proc Natl Acad Sci U S A. 1992;89(14):6580–4. doi: 10.1073/pnas.89.14.6580. PubMed PMID: 1631159; PubMed Central PMCID: PMCPMC49545.

18. Symons J, Chopra A, Malatinkova E, De Spiegelaere W, Leary S, Cooper D, et al. HIV integration sites in latently infected cell lines: evidence of ongoing replication. Retrovirology. 2017;14(1):2. Epub 20170113. doi: 10.1186/s12977-016-0325-2. PubMed PMID: 28086908; PubMed Central PMCID: PMCPMC5237276.

19. Kopera HC, Larson PA, Moldovan JB, Richardson SR, Liu Y, Moran JV. LINE-1 Cultured Cell Retrotransposition Assay. Methods Mol Biol. 2016;1400:139–56. doi: 10.1007/978-1-4939-3372-3_10. PubMed PMID: 26895052; PubMed Central PMCID: PMCPMC5070806.

20. Sloan RD, Kuhl BD, Mesplède T, Münch J, Donahue DA, Wainberg MA. Productive Entry of HIV-1 during Cell-to-Cell Transmission via Dynamin-Dependent Endocytosis. J Virol. 2013;87(14):8110–23. doi: 10.1128/Jvi.00815-13. PubMed PMID: WOS:000321017700030.

21. Speck RR, Flexner C, Tian CJ, Fu XF. Comparison of human immunodeficiency virus type 1 Pr55 and Pr160 processing intermediates that accumulate in primary and transformed cells treated with peptidic and nonpeptidic protease inhibitors. Antimicrob Agents Ch. 2000;44(5):1397–403. doi: Doi 10.1128/Aac.44.5.1397-1403.2000. PubMed PMID: WOS:000086625400052.

22. Furman PA, Fyfe JA, Stclair MH, Weinhold K, Rideout JL, Freeman GA, et al. Phosphorylation of 3’-Azido-3’-Deoxythymidine and Selective Interaction of the 5’-Triphosphate with Human-Immunodeficiency-Virus Reverse-Transcriptase. P Natl Acad Sci USA. 1986;83(21):8333–7. doi: DOI 10.1073/pnas.83.21.8333. PubMed PMID: WOS:A1986E643100065.

23. De Meyer S, Azijn H, Surleraux D, Jochmans D, Tahri A, Pauwels R, et al. TMC114, a novel human immunodeficiency virus type 1 protease inhibitor active against protease inhibitor-resistant viruses, including a broad range of clinical isolates. Antimicrob Agents Ch. 2005;49(6):2314–21. doi: 10.1128/Aac.49.6.2314-2321.2005. PubMed PMID: WOS:000229521600020.

24. Hsieh SH, Yu FH, Huang KJ, Wang CT. HIV-1 reverse transcriptase stability correlates with Gag cleavage efficiency: reverse transcriptase interaction implications for modulating protease activation. J Virol. 2023;97(9). doi: ARTN e00948-23 10.1128/jvi.00948-23. PubMed PMID: WOS:001167283600025.

25. Hill MK, Hooker CW, Harrich D, Crowe SM, Mak J. Gag-Pol supplied in is efficiently packaged and supports viral function in human immunodeficiency virus type 1. J Virol. 2001;75(15):6835–40. doi: Doi 10.1128/Jvi.75.15.6835-6840.2001. PubMed PMID: WOS:000169870700011.

26. Shimura K, Kodama E, Sakagami Y, Matsuzaki Y, Watanabe W, Yamataka K, et al. Broad Antiretroviral activity and resistance profile of the novel human immunodeficiency virus integrase inhibitor elvitegravir (JTK-303/GS-9137). J Virol. 2008;82(2):764–74. doi: 10.1128/Jvi.01534-07. PubMed PMID: WOS:000252229100018.

27. Hightower KE, Wang RL, DeAnda F, Johns BA, Weaver K, Shen YN, et al. Dolutegravir (S/GSK1349572) Exhibits Significantly Slower Dissociation than Raltegravir and Elvitegravir from Wild-Type and Integrase Inhibitor-Resistant HIV-1 Integrase-DNA Complexes. Antimicrob Agents Ch. 2011;55(10):4552–9. doi: 10.1128/Aac.00157-11. PubMed PMID: WOS:000294952600009.

28. D’Souza V, Summers MF. How retroviruses select their genomes. Nat Rev Microbiol. 2005;3(8):643–55. doi: 10.1038/nrmicro1210. PubMed PMID: WOS:000230879700015.

29. Llewellyn GN, Hogue IB, Grover JR, Ono A. Nucleocapsid Promotes Localization of HIV-1 Gag to Uropods That Participate in Virological Synapses between T Cells. Plos Pathog. 2010;6(10). doi: ARTN e1001167 10.1371/journal.ppat.1001167. PubMed PMID: WOS:000283652200039.

30. Ono A, Ablan SD, Lockett SJ, Nagashima K, Freed EO. Phosphatidylinositol (4,5) bisphosphate regulates HIV-1 gag targeting to the plasma membrane. Mol Biol Cell. 2004;15:122a–3a. PubMed PMID: WOS:000224648801025.

31. Freed EO, Orenstein JM, Bucklerwhite AJ, Martin MA. Single Amino-Acid Changes in the Human-Immunodeficiency-Virus Type-1 Matrix Protein Block Virus Particle-Production. J Virol. 1994;68(8):5311–20. doi: Doi 10.1128/Jvi.68.8.5311-5320.1994. PubMed PMID: WOS:A1994NW97800072.

32. Freed EO, Englund G, Martin MA. Role of the Basic Domain of Human-Immunodeficiency-Virus Type-1 Matrix in Macrophage Infection. J Virol. 1995;69(6):3949–54. doi: Doi 10.1128/Jvi.69.6.3949-3954.1995. PubMed PMID: WOS:A1995QX93100090.

33. Chukkapalli V, Oh SJ, Ono A. Opposing mechanisms involving RNA and lipids regulate HIV-1 Gag membrane binding through the highly basic region of the matrix domain. P Natl Acad Sci USA. 2010;107(4):1600–5. doi: 10.1073/pnas.0908661107. PubMed PMID: WOS:000273974600067.

34. Monde K, Chukkapalli V, Ono A. Assembly and replication of HIV-1 in T cells with low levels of phosphatidylinositol-(4,5)-bisphosphate. J Virol. 2011;85(7):3584–95. Epub 20110126. doi: 10.1128/JVI.02266-10. PubMed PMID: 21270152; PubMed Central PMCID: PMCPMC3067840.

35. Joshi A, Nagashima K, Freed EO. Mutation of dileucine-like motifs in the human immunodeficiency virus type 1 capsid disrupts virus assembly, Gag-Gag interactions, Gag-membrane binding, and virion maturation. J Virol. 2006;80(16):7939–51. doi: 10.1128/Jvi.00355-06. PubMed PMID: WOS:000239557700016.

36. von Schwedler UK, Stray KM, Garrus JE, Sundquist WI. Functional surfaces of the human immunodeficiency virus type 1 capsid protein. J Virol. 2003;77(9):5439–50. doi: 10.1128/Jvi.77.9.5439-5450.2003. PubMed PMID: WOS:000182297200041.

37. Pornillos O, Ganser-Pornillos BK, Kelly BN, Hua YZ, Whitby FG, Stout CD, et al. X-Ray Structures of the Hexameric Building Block of the HIV Capsid. Cell. 2009;137(7):1282–92. doi: 10.1016/j.cell.2009.04.063. PubMed PMID: WOS:000267373400021.

38. Shin R, Tzou YM, Krishna NR. Structure of a Monomeric Mutant of the HIV-1 Capsid Protein. Biochemistry-Us. 2011;50(44):9457–67. doi: 10.1021/bi2011493. PubMed PMID: WOS:000296304200005.

39. Sherer NM, Lehmann MJ, Jimenez-Soto LF, Ingmundson A, Horner SM, Cicchetti G, et al. Visualization of retroviral replication in living cells reveals budding into multivesicular bodies. Traffic. 2003;4(11):785–801. doi: DOI 10.1034/j.1600-0854.2003.00135.x. PubMed PMID: WOS:000185876900007.

40. McCluskey A, Daniel JA, Hadzic G, Chau N, Clayton EL, Mariana A, et al. Building a Better Dynasore: The Dyngo Compounds Potently Inhibit Dynamin and Endocytosis. Traffic. 2013;14(12):1272–89. doi: 10.1111/tra.12119. PubMed PMID: WOS:000339890000001.

41. Gratton S, Cheynier R, Dumaurier MJ, Oksenhendler E, Wain-Hobson S. Highly restricted spread of HIV-1 and multiply infected cells within splenic germinal centers. P Natl Acad Sci USA. 2000;97(26):14566–71. doi: DOI 10.1073/pnas.97.26.14566. PubMed PMID: WOS:000165993700104.

42. Jung A, Maier R, Vartanian JP, Bocharov G, Jung V, Fischer U, et al. Recombination - Multiply infected spleen cells in HIV patients. Nature. 2002;418(6894):144–. doi: 10.1038/418144a. PubMed PMID: WOS:000176710400029.

43. Josefsson L, Palmer S, Faria NR, Lemey P, Casazza J, Ambrozak D, et al. Single Cell Analysis of Lymph Node Tissue from HIV-1 Infected Patients Reveals that the Majority of CD4 T-cells Contain One HIV-1 DNA Molecule. Plos Pathog. 2013;9(6). doi: ARTN e1003432 10.1371/journal.ppat.1003432. PubMed PMID: WOS:000321206600039.

44. Welsch S, Keppler OT, Habermann A, Allespach I, Krijnse-Locker J, Kräusslich HG. HIV-1 buds predominantly at the plasma membrane of primary human macrophages. Plos Pathog. 2007;3(3). doi: ARTN e36 10.1371/journal.ppat.0030036. PubMed PMID: WOS:000248495200013.

45. Grigorov B, Arcanger F, Roingeard P, Darlix JL, Muriaux D. Assembly of infectious HIV-1 in human epithelial and T-lymphoblastic cell lines. J Mol Biol. 2006;359(4):848–62. doi: 10.1016/j.jmb.2006.04.017. PubMed PMID: WOS:000238682800003.

46. Daecke J, Fackler OT, Dittmar MT, Kräusslich HG. Involvement of clathrin-mediated endocytosis in human immunodeficiency virus type 1 entry. J Virol. 2005;79(3):1581–94. doi: 10.1128/Jvi.79.3.1581-1594.2005. PubMed PMID: WOS:000226634300025.

47. Miyauchi K, Kim Y, Latinovic O, Morozov V, Melikyan GB. HIV Enters Cells via Endocytosis and Dynamin-Dependent Fusion with Endosomes. Cell. 2009;137(3):433–44. doi: 10.1016/j.cell.2009.02.046. PubMed PMID: WOS:000265677200014.

48. Padilla-Parra S, Marin M, Gahlaut N, Suter R, Kondo N, Melikyan GB. Fusion of Mature HIV-1 Particles Leads to Complete Release of a Gag-GFP-Based Content Marker and Raises the Intraviral pH. Plos One. 2013;8(8). doi: ARTN e71002 10.1371/journal.pone.0071002. PubMed PMID: WOS:000326473200030.

49. Rodari A, Darcis G, Van Lint CM. The Current Status of Latency Reversing Agents for HIV-1 Remission. Annu Rev Virol. 2021;8(1):491–514. doi: 10.1146/annurev-virology-091919-103029. PubMed PMID: 34586875.

50. Ahlenstiel CL, Symonds G, Kent SJ, Kelleher AD. Block and Lock HIV Cure Strategies to Control the Latent Reservoir. Front Cell Infect Microbiol. 2020;10:424. Epub 20200814. doi: 10.3389/fcimb.2020.00424. PubMed PMID: 32923412; PubMed Central PMCID: PMCPMC7457024.

51. Xie Y, Rosser JM, Thompson TL, Boeke JD, An WF. Characterization of L1 retrotransposition with high-throughput dual-luciferase assays. Nucleic Acids Res. 2011;39(3). doi: ARTN e16 10.1093/nar/gkq1076. PubMed PMID: WOS:000287257500005.

52. Huang MJ, Orenstein JM, Martin MA, Freed EO. P6(Gag) Is Required for Particle-Production from Full-Length Human-Immunodeficiency-Virus Type-1 Molecular Clones Expressing Protease. J Virol. 1995;69(11):6810–8. doi: Doi 10.1128/Jvi.69.11.6810-6818.1995. PubMed PMID: WOS:A1995RZ10000025.

53. Ott DE, Coren LV, Chertova EN, Gagliardi TD, Nagashima K, Sowder RC, 2nd, et al. Elimination of protease activity restores efficient virion production to a human immunodeficiency virus type 1 nucleocapsid deletion mutant. J Virol. 2003;77(10):5547–56. doi: 10.1128/jvi.77.10.5547-5556.2003. PubMed PMID: 12719547; PubMed Central PMCID: PMCPMC154014.

54. Sasaki H, Arai H, Kikuchi E, Saito H, Seki K, Matsui T. Novel electron microscopic staining method using traditional dye, hematoxylin. Sci Rep-Uk. 2022;12(1). doi: ARTN 7756 10.1038/s41598-022-11523-y. PubMed PMID: WOS:000796701700015.

55. Satou Y, Katsuya H, Fukuda A, Misawa N, Ito J, Uchiyama Y, et al. Dynamics and mechanisms of clonal expansion of HIV-1-infected cells in a humanized mouse model. Sci Rep. 2017;7(1):6913. Epub 20170731. doi: 10.1038/s41598-017-07307-4. PubMed PMID: 28761140; PubMed Central PMCID: PMCPMC5537293.

56. Li H, Durbin R. Fast and accurate short read alignment with Burrows-Wheeler transform. Bioinformatics. 2009;25(14):1754–60. doi: 10.1093/bioinformatics/btp324. PubMed PMID: WOS:000267665900006.

57. Broad I. Picard toolkit (software). GitHub repository2019.

